# DSA-DeepFM: A Dual-Stage Attention-Enhanced DeepFM Model for Predicting Anticancer Synergistic Drug Combinations

**DOI:** 10.1101/2024.11.02.621696

**Authors:** Yuexi Gu, Yongheng Sun, Louxin Zhang, Jian Zu

**Affiliations:** School of Mathematics and Statistics, Xi’an Jiaotong University, 710049, Shaanxi, People’s Republic of China; Department of Mathematics and Centre for Data Science and Machine Learning, National University of Singapore, Singapore 119076, Singapore

**Keywords:** Drug combination and synergy, factorization machine, dual-stage attention, deep learning, AI

## Abstract

**Motivation:** Drug combinations are crucial in combating drug resistance, reducing toxicity, and improving therapeutic outcomes in the treatment of complex diseases. As the number of available drugs grows, the potential combinations increase exponentially, making it impractical to rely solely on biological experiments to identify synergistic drug pairs. Consequently, machine learning methods are increasingly used to systematically screen for synergistic drug combinations. However, most current approaches prioritize predictive performance by integrating auxiliary information or increasing model complexity. By overlooking the biological mechanisms behind feature interactions, their effectiveness in predicting drug synergy can be limited.

**Results:** We present DSA-DeepFM, a deep learning model that integrates a dual-stage attention (DSA) mechanism with Factorization Machines (FMs) to improve drug synergy prediction by addressing complex biological feature interactions. The model incorporates categorical and auxiliary numerical inputs, embedding them into high-dimensional spaces and then fusing them through the DSA mechanism to capture both field-aware and embedding-aware patterns. These patterns are then processed by a DeepFM module, which captures low-order and high-order feature interactions before making final predictions. Validation testing demonstrates that DSA-DeepFM significantly outperforms traditional machine learning and state-of-the-art deep learning models. Additionally, t-SNE visualizations confirm the model’s discriminative power at various stages. As a case study, we use our model to identify eight novel synergistic drug combinations, three of which are well-supported by existing wet-lab experiments, underscoring its practical utility and potential for future applications.

---

Compared to single-agent therapy, combination therapy utilizes multiple drugs targeting different biological pathways and mechanisms in the treatment of complex diseases such as cancer, HIV, and diabetes [Bar-On et al., 2018, Coates et al., 2020, Fisusi and Akala, 2019, Jaaks et al., 2022, Rea et al., 2018, Tan et al., 2012, Zhao et al., 2013]. Drug combinations can improve efficacy while minimizing the toxic side effects associated often with monotherapy. For example, Nussinov et al. [2013] proposed the ‘Pathway Drug Cocktails’ strategy for treating tumors with Ras mutations. This approach facilitates the action of multiple drugs on various cellular processes involved in tumorigenesis. Moreover, drug combinations are very useful in mitigating drug resistance. For example, combining epidermal growth factor receptor tyrosine kinase inhibitor with various drugs is investigated as an anti-resistance strategy for treating lung cancer [Tong et al., 2017].

Nevertheless, inappropriate drug combinations can lead to antagonistic effects or even worsen disease progression, as evidenced by a clinical study involving 48 patients diagnosed with idiosyncratic drug-induced liver injury who were taking two drugs [Benesic et al., 2019]. Therefore, accurately designing synergistic combination regimens is essential for optimizing therapeutic outcomes in precision medicine.

The increasing number of available drugs poses a great challenge for the screening of synergistic combinations through clinical experiments. Advances in high-throughput drug screening have enabled large-scale drug combination studies [Al-Lazikani et al., 2012], resulting in the creation of several notable databases containing synergistic and antagonistic interactions on different cell lines, such as O’Neil’s dataset [O’Neil et al., 2016], the NCI-A Large Matrix of Anti-Neoplastic Agent Combinations (ALMANAC) dataset [Holbeck et al., 2017], and DrugCombDB [Liu et al., 2020, Zheng et al., 2021].

This facilitates the development of scalable AI and machine learning approaches for efficiently identifying synergistic drug combinations, complementing conventional experimental methods ([Abbasi and Rousu, 2024, Besharatifard and Vafaee, 2024] for review). In particular, recent advancements in deep learning have significantly improved the prediction of drug combination effects [Alam et al., 2024, Bertin et al., 2023, Hu et al., 2023, Jiang et al., 2020, Kuru et al., 2021, Liu et al., 2022, Monem et al., 2023, Sun et al., 2020, Torkamannia et al., 2023, Tsepa et al., 2023, Wang et al., 2022a,b, Xu et al., 2023, Zhang et al., 2022].

Although these models effectively predict drug synergies, significant challenges remain [Abbasi and Rousu, 2024, Besharatifard and Vafaee, 2024]. For instance, some models incorporate diverse drug and the cell line features to enhance performance; however, this increases computational complexity and introduces noise or irrelevant data, which can reduce overall effectiveness. Furthermore, merging data from multiple databases often results in incomplete entries, leading to under-utilization of available data. Models that incorporate pre-trained language models to improve feature representation may also struggle to capture high-order interactions between different feature types, thus limiting predictive capability.

We propose an end-to-end Dual-Stage Attention-enhanced Deep Factorization Machine (DSA-DeepFM) model for predicting synergistic drug combinations as a classification task. It integrates categorical and auxiliary numerical profiles of drugs and cell line to enrich feature representation. Unlike traditional DeepFM models used in other domains [Gao and Wu, 2022, Guo et al., 2017, Su et al., 2022], DSA-DeepFM uses a DSA mechanism to fuse profiles in embedding and field spaces, improving feature learning without increasing dimensionality. Its FM module captures importance of features and their interactions [Rendle, 2010], while its Deep Neural Network (DNN) module captures higher-order interactions. A residual connection to the prediction module helps to maintain gradients during training, preventing vanishing gradients and supporting more effective learning.

## Materials and Methods

### Datasets

The databases used in this study are summarized in Table 1.

**Table 1.**
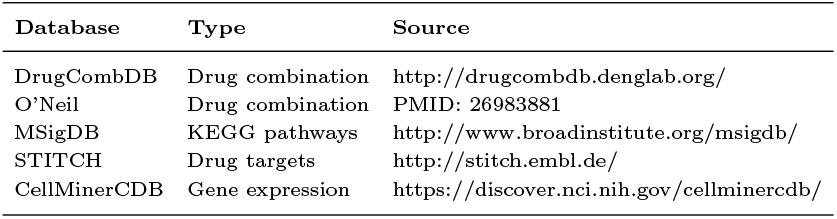
A summary of the databases used in this study.

#### Drug combination datasets

DrugCombDB (accessed in Feb 2023; Liu et al. [2020]) includes 448,555 drug combinations, involving 2,887 specific drugs and 124 human cancer cell lines across 14 tissue types. This database integrates data from various sources, including NCI-ALMANAC project [Holbeck et al., 2017], the AstraZeneca–Sanger dataset [Menden et al., 2019], manually curated data from the literature, DCDB [Liu et al., 2014] and the FDA Orange Book [Food and Drug Administration, 2005].

The O’Neil (also called Merck) dataset comprises 22,737 results from experiments across 39 cancer cell lines and 38 anticancer drugs (accessed in Feb 2023, O’Neil et al. [2016]).

#### Drug molecular fingerprints

The Simplified Molecular Input Line Entry System (SMILES) representations of drugs were downloaded from the DrugCombDB database.

#### KEGG pathways

186 KEGG pathways, involving 5,244 genes, were downloaded from the Molecular Signatures Database (MSigDB) (C2 collection, accessed in Feb 2023, Liberzon et al. [2015]).

#### Drug target data

We downloaded the drug target data from the Search Tool for Interactions of Chemicals (STITCH) database [Szklarczyk et al., 2016]. This database aggregates drug-protein interactions from multiple databases and literature sources.

#### Cell line profiles

RNA-Seq gene expression data for the cell lines in DrugCombDB were obtained from the CellMiner Cross Database (CellMinerCDB) [Rajapakse et al., 2018, Luna et al., 2021]. This platform integrates multiple cancer cell line datasets, including NCI-60, the Broad Cancer Cell Line Encyclopedia, and Sanger/MGH Genomics of Drug Sensitivity in Cancer.

## Methods

### Data preprocessing

Following Preuer et al. [2018], we labeled drug pairs with synergy scores greater than 30 as positive (synergistic), and those with scores below 0 as negative (antagonistic) for the O’Neil dataset.

In cases where a drug combination has multiple entries with varying synergy outcomes across the two databases, majority voting was employed to determine the overall synergistic property of the combination.

We converted the SMILES representations of drugs into the 1024-bit extended connectivity fingerprints with a diameter of 6 (ECFP6) using the Morgan algorithm in RDKit [Landrum et al., 2013]. The Tanimoto coefficient was then calculated based on the ECFP6 fingerprints.

To standardize the RNA-sequencing gene expression data from CellMinerCDB, we applied z-score normalization.

### Validation testing

To assess our model’s performance, we compared it against nine state-of-the-art prediction methods using five-fold cross-validation, with the data split into training, validation, and test sets in a 3:1:1 ratio.

We also conduct validation tests for two more challenging leave-cell line or tissue-out cases.

- Leave-cell line-out: Cell lines in the test set were not present in the training set.
- Leave-tissue-out: The cell lines in a tissue appearing in the test set were not included in the training set.

For the leave-out scenarios, we employed the same cross-validation to compare the prediction performance of DSA-DeepFM against other models. Since O’Neil dataset is small, it is hard to design proper tests for these tests and thus we only validated the methods on DrugCombDB.

Eight metrics were used to evaluate different models: area under the receiver operating characteristic curve (AUC-ROC), area under the precision-recall curve (AUC-PR), accuracy (ACC), precision, recall, F1 score, Cohen’s kappa (Kappa), and balanced accuracy (BACC). (For their definitions, see Section 2 of the Supplementary Document).

### Configuring the model’s hyperparameters

The key hyperparameters of the model were tuned based on performance on the validation datasets, measured by AUC.

#### The size of the FM module

The embedding layer of FM converts high-dimensional categorical data into dense vectors. Testing embedding dimensions from 64 to 1024 for the module showed consistent performance improvements as the dimension increased (Figure 1a). Performance stabilized at 512, with both mean and variance of AUC closely matching those at 1024. Therefore, 512 was chosen as the size of FM.

**Fig. 1.**
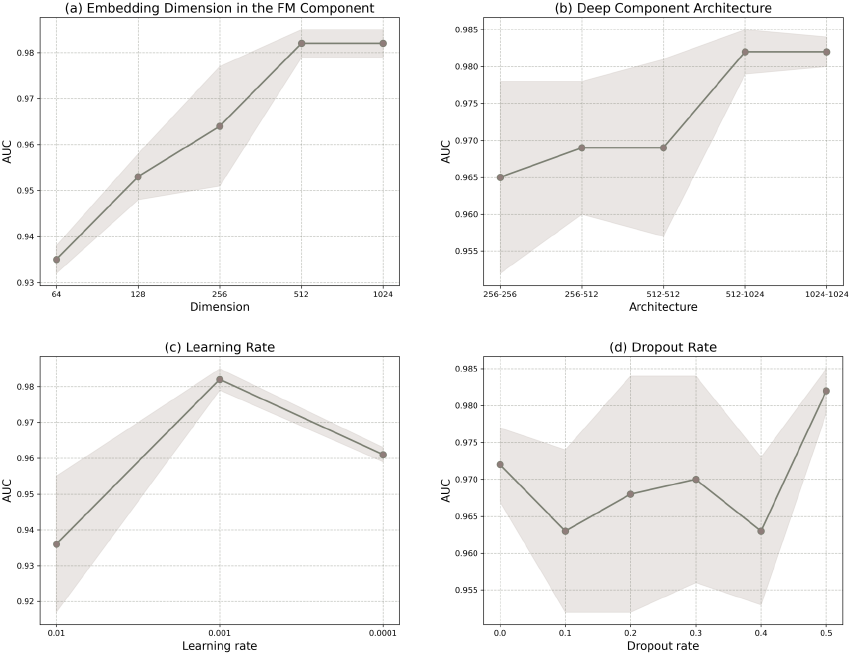
Parameter sensitivity analysis. (a) AUC vs. the dimension of the FM component. (b) AUC vs. the size of the deep component. (c) AUC vs. the initial learning rate during training. (d) AUC vs. the dropout rate in the hidden layers. The shaded area represents the variation of the AUC.

#### The size of the DNN module

The module consists of a two-layer fully connected network. Therefore, various combinations of layer dimensions were tested to optimize performance (Figure 1b, Section 1.1 of the Supplementary Document). Starting with a 256 × 256 network, increasing the second layer’s size improved the mean AUC and reduced variance. Further increases in the first and second layers yielded diminishing returns in performance. Given that the 512 × 1024 and 1024 × 1024 configurations produced similar results, the 512 × 1024 network was selected to balance performance and model complexity.

#### Learning rate

The model was trained using the Adam optimizer with learning rate decay. After tuning, a learning rate of 0.001 was selected, as it significantly improved mean AUC and reduced variance compared to 0.01, while a further decrease to 0.0001 caused a decline in performance (Figure 1c, Section 1.2 of the Supplementary Document).

#### Dropout rate

We tested dropout rates ranging from 0.0 to 0.5 to prevent overfitting. AUC increased gradually with dropout rates between 0.1 and 0.3, declined at 0.4 and then peaked at 0.5 (Figure 1d, Section 1.3 of the Supplementary Document). Overall, dropout had minimal impact on the model’s performance. We set the drop rate to 0.5 for our model.

### Ablation analysis

The following five variants of the DSA-DeepFM were designed to isolate the effects of different mechanisms (data types, attention, residuals) on the model’s performance:

- **DSA-DeepFM-cat**: Use only categorical data and handles inputs via embedding and feeds them into both FM and DNN components.
- **DSA-DeepFM-aux**: Processes only auxiliary data through a two-layer network before inputting it into FM and DNN.
- **DSA-DeepFM-w/o-attn**: This version processes both categorical and auxiliary data, but instead of using attention, it concatenates the vectors before inputting them into FM and DNN.
- **DSA-DeepFM-w/o-res**: In this version, residual connectivity is removed from the prediction stage, and predictions are made using a three-layer fully connected network.
- **DSA-DeepFM-attn-rev**: Here, the attention mechanism is applied in reverse order. Attention scores are first calculated in the 512-dimensional embedding space, followed by the 3-dimensional field space.

## Results

### Deep Learning Model

The proposed model is illustrated in Figure 2 and implemented using the TensorFlow backend (Version 1.14.0). The input to the model consists of two types of data: the auxiliary numerical data, derived from the molecular profiles of drugs and the cell lines, and the one-hot encoded categorical data.

**Fig. 2.**
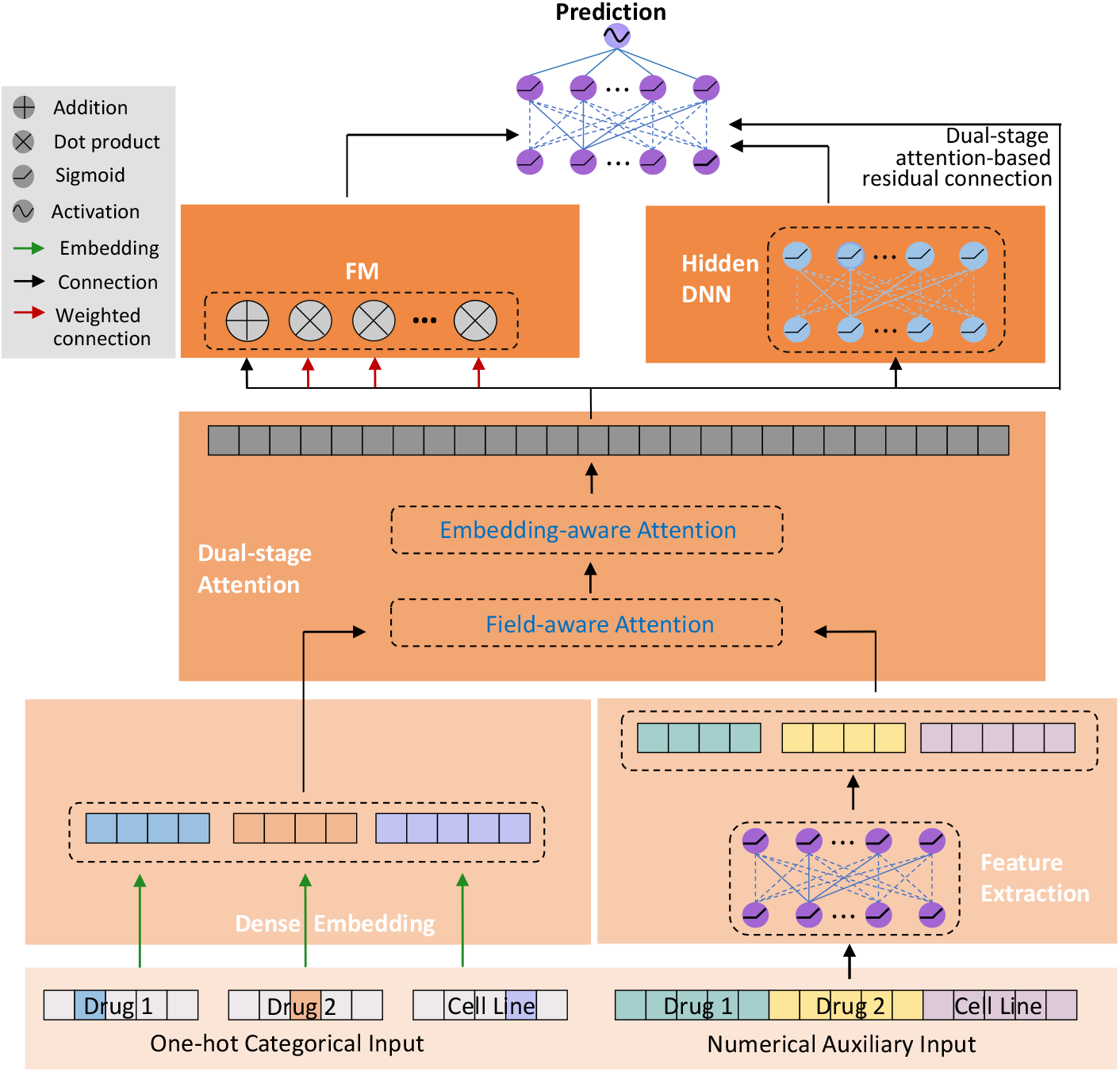
The architecture of DSA-DeepFM. The input consists of two types of data: Drug and cell line names (one-hot encoded) and auxiliary features such as drug molecular fingerprints (FPs) and cell line gene expression (GE) data. The Dual-stage Attention module allows both feature types to learn field-shared and embedding-space-shared representations while preserving their unique characteristics. The concatenated vector is then fed into both the FM layer and the Hidden DNN module. FM captures low-order feature interactions, while Hidden DNN models high-order interactions. Finally, the combined features are passed to the prediction module, which uses a residual structure to improve the prediction of synergistic drug combinations.

The categorical input is mapped to a high-dimensional feature space by the dense embedding layer, enabling the model to learn useful latent features during training. Simultaneously, a feature extraction layer processes the auxiliary information to identify significant features. These extracted features are then fused with the learned representations from the embedding layer using the DSA mechanism. This integration enhances the model’s ability to classify and characterize features by effectively combining known auxiliary information with the latent representations.

Next, the embedding features and auxiliary features are concatenated into a single input vector, which is shared between the FM layer [Rendle, 2010] and the Hidden DNN module. FM attempts to capture low-order feature interactions, while Hidden DNN learns high-order feature interactions. The low-order and high-order interactions are then concatenated and fed to the prediction module, along with the output from the DSA module, where residual connections based on the DSA mechanism are adopted to mitigate the vanishing gradient problem.

#### Feature generation

##### Dense Embedding layer for the categorical input

Assume there are *m* drugs and *n* cell lines. We use 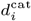 and 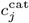 to denote the categorical data on the *i*-th drug and the *j*-th cell line, respectively, for each *i* and *j*. For each pair of drugs and each cell line, the categorical input is the triplet 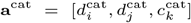. The embedding layer of our model maps each input triplet to a dense numeric vector in ℝ^3*×E*^ using an embedding dictionary [Guo et al., 2017, Gao and Wu, 2022]. Here, each 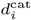 is mapped to 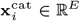 and each 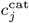 is mapped to 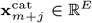. (The embedding size *E* was set to 512 in our study.) The dictionary 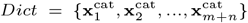 contains the trainable latent vectors, which can be interpreted as the weights of a fully connected network and will be learned during the training process [Su et al., 2022].

##### Feature Extraction module for the auxiliary feature input

Let 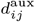 be the Tanimoto coefficient between the ECFP6 fingerprints of drug *d*_*i*_ and *d*_*j*_. For each drug *d*_*i*_, its auxiliary information is represented as 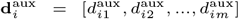.

Similarly, for each *k*, the auxiliary information of cell line *c*_*k*_ is 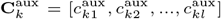, where *l* is the number of genes used in our model, and 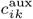 represents the normalized RNA-seq value of the *k*-th gene in the cell line. For each pair of drugs and each cell line, the corresponding feature input is 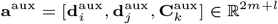.

To align the auxiliary information with the categorical embedding, we implemented a two-layer fully connected network to extract latent features from the feature input triplets, which maps each **a**^aux^ to **x**^aux^ in the same feature space ℝ^3*×E*^ as the categorical embedding vectors. The extracted numerical feature is denoted 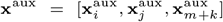, where 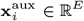.

#### The DSA module

To integrate the categorical embedding and numerical features, we propose a DSA mechanism (Figure 3). The inputs to this module are the categorical embedding vector 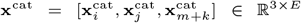 and the feature vector 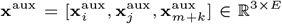. The DSA module has two components: Field-aware attention (Figure 3a) and embedding-aware attention (Figure 3b).

**Fig. 3.**
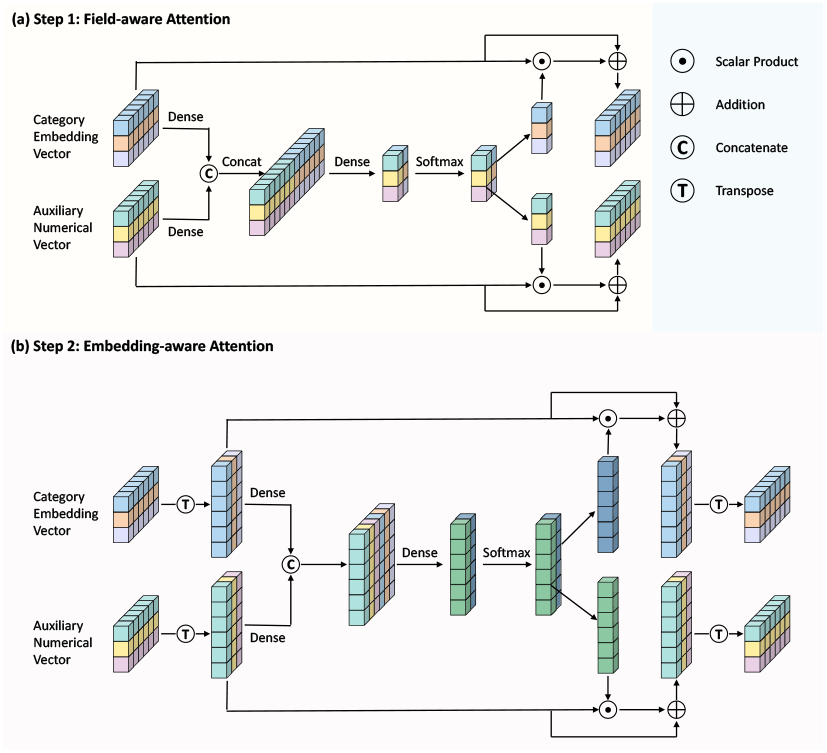
The dual-stage attention module. (a) Field-aware Attention. It computes the shared patterns in the embedding space along with the attention scores for each field. (b) Embedding-aware Attention. It extracts the shared patterns in the field space and the attention scores for each embedding dimension.

##### Field-aware attention module

In this stage, attention scores are computed for each row (corresponding to a field) in the feature matrices. The categorical embedding vectors 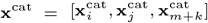 and the auxiliary numerical vectors 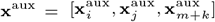 are first updated using two different dense operations as:

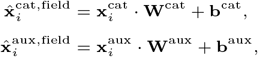

where **W**^cat^, **W**^aux^ ∈ ℝ ^*E×E*^ are the weight matrices, and **b**^cat^, **b**^aux^ ∈ ℝ ^*E*^ are the corresponding bias terms.

These updated vectors are concatenated into a single vector, which is then passed through another dense layer with weight matrix **W**^field^ ∈ ℝ ^2*E×*2^ and bias **b**^field^ ∈ ℝ ^2^. This operation maps the features into vectors in the same embedding space, enhancing learning shared patterns. A softmax transformation is then applied to calculate the attention scores for the *i*-th field:

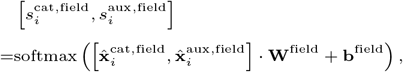

which are used to assign importance to each field to scale the original vectors. Precisely, the final field-aware categorical vector **x**^cat,field^ and auxiliary vector **x**^aux,field^ are obtained via a residual connection as:

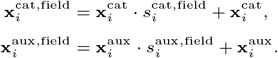

##### Embedding-aware attention module

In this component, attention scores will be computed for each feature dimension within the field space. Both the field-aware categorical vectors 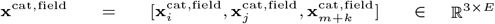 and field-aware auxiliary vectors 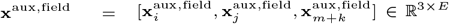 are first transposed so that each row now corresponds to an embedding dimension, and each column corresponds to a field.

Similar to the field-aware attention, the two vectors are passed through two dense operations and concatenated to merge their vectors into the field space. Assume the dense weight matrices are **W**^cat,emb^, **W**^aux,emb^ ∈ ℝ^3*×*3^, and assume the corresponding bias terms used are **b**^cat,emb^, **b**^aux,emb^ ∈ ℝ^3^. The updated categorical vectors and auxiliary vectors for the *j*-th embedding dimension are:

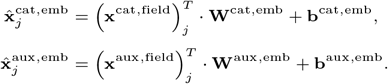

These vectors 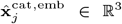 and 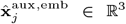 are then concatenated into a single vector and are fed to another dense layer with weight matrix **W**^emb^ ∈ ℝ^6*×*2^ and bias **b**^emb^ ∈ ℝ^2^. The attention score for the *j*-th embedding dimension can be calculated as:

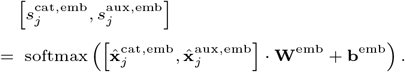

The embedding-aware categorical vectors **x**^cat,emb^ and auxiliary vectors **x**^aux,emb^ are further updated as:

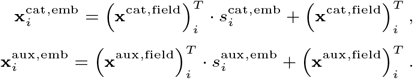

Finally, the vectors (**x**^cat,emb^)^*T*^ and (**x**^aux,emb^)^*T*^ are flatten and concatenated to form the final feature vector **X**:

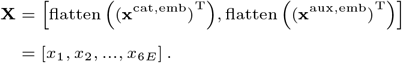

**X** will be the input to both the FM layer and DNN module.

#### FM layer

The FM mechanism, initially introduced for extracting interaction features in recommender system design [Rendle, 2010], captures feature relationships through the inner product of two latent features. We use FM to assess both the importance of the features and the influence of their interactions for the input vector **X** = [*x*_1_, *x*_2_, …, *x*_6*E*_]. The output of FM is:

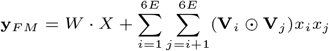

where · means matrix multiplication and ⊙ means element-wise multiplication, *W* ∈ ℝ^*K×*6*E*^ and **V**_*i*_ ∈ ℝ^*K*^ are the trainable parameters, and the hyper-parameter *K* was set to 1024 in this study. Considering *w*_*i*_ as the weight of *i*-th feature, we use 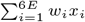 to represent the feature importance. Furthermore, we use ⟨**V**_*i*_, **V**_*j*_ ⟩ to capture the interaction between the features *x*_*i*_ and *x*_*j*_. As a result, 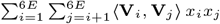 represents the overall influence of the pairwise feature interactions.

#### Hidden DNN module

It is the ‘deep’ part of our model. After the FM layer had learnt the importance of features and their pairwise interactions, DNN Hidden is used to learn the higher-order interaction among the features. This module is a two-layer Feed-Forward neural network. Its input vector is **X**^0^ = [*x*_1_, *x*_2_, …, *x*_6*E*_] computed in the dual-state attention component and its output **y**_*DNN*_ is:

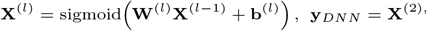

where **W**^(*l*)^ and **b**^(*l*)^ are the learnable weight and bias parameters for the *l*-th layer.

#### Prediction Module

FM and Hidden DNN learn the latent features and their high-order interactions, respectively. Lastly, we concatenate these two features as the input to the prediction module. Additionally, a DSA based residual connection is added to enhance the performance of our model. The input is denoted as 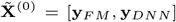. Firstly, we map 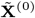 into the same feature space with **X**^(0)^ to align the dimension of these two vectors, resulting in:

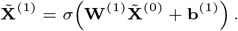

Next, a residual connection based on the DSA mechanism is introduced to combine 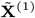 with the original raw feature **X**^(0)^, improving feature representation and mitigating the vanishing gradient problem.

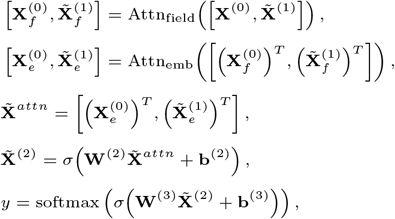

where Attn_field_ and Attn_emb_ denote field-aware and embedding-aware output from the DSA module, respectively. The output *y* represents the probability that the triplet [drug 1, drug 2, cell line] exhibits a synergistic effect.

### Ablation analysis

#### The impact of different mechanisms on model performance

Ablation experiments were conducted by applying different variants of DSA-DeepFM (which are described in the Method section) on the DrugCombDB dataset to investigate the following questions:

- Does the use of both categorical and auxiliary data improve performance compared to using either type of data alone?
- Does incorporating an attention mechanism enhance model performance?
- When using an attention mechanism, which architecture performs better: starting with the field space of dimension three followed by the embedding space of dimension 512, or the reverse?
- Does adding a residual mechanism to the prediction module improve performance?

Since the residual structure in DSA-DeepFM is based on a DSA mechanism, we excluded the residual connectivity for a fair comparison across all the models listed above. Here, we focus on AUC, as the models had similar performance across different metrics.

Table 2 indicates that DSA-DeepFM outperformed all variants, achieving a mean AUC of 0.982 with a standard deviation of 0.003. Comparing DSA-DeepFM-cat, DSA-DeepFM-aux and DSA-DeepFM-w/o-attn reveals that the best results were achieved when only categorical inputs were used, with a mean AUC of 0.897 (±0.004). This suggests that the feature fusion mechanism significantly improves the performance of a model, and the embedding layer effectively captures essential hidden features from sparse inputs.

**Table 2.** Results of the ablation study on the DrugCombDB Dataset.

| Model | AUC-ROC | ACC | Precision | Recall | F1 | AUC-PR | Kappa | BACC |
| --- | --- | --- | --- | --- | --- | --- | --- | --- |
| DSA-DeepFM | **0.982**$\pm 0.003$ | **0.962**$\pm 0.004$ | **0.943**$\pm 0.006$ | **0.932**$\pm 0.011$ | **0.937**$\pm 0.008$ | **0.968**$\pm 0.005$ | **0.910**$\pm 0.011$ | **0.954**$\pm 0.006$ |
| DSA-DeepFM-cat | 0.897$\pm 0.004$ | 0.850$\pm 0.002$ | 0.813$\pm 0.006$ | 0.656$\pm 0.006$ | 0.726$\pm 0.006$ | 0.832$\pm 0.006$ | 0.624$\pm 0.007$ | 0.795$\pm 0.003$ |
| DSA-DeepFM-aux | 0.825$\pm 0.003$ | 0.798$\pm 0.002$ | 0.749$\pm 0.006$ | 0.504$\pm 0.006$ | 0.602$\pm 0.004$ | 0.718$\pm 0.004$ | 0.474$\pm 0.005$ | 0.715$\pm 0.003$ |
| DSA-DeepFM-w/o-attn | 0.890$\pm 0.003$ | 0.843$\pm 0.002$ | 0.801$\pm 0.006$ | 0.643$\pm 0.007$ | 0.713$\pm 0.004$ | 0.820$\pm 0.005$ | 0.607$\pm 0.005$ | 0.787$\pm 0.003$ |
| DSA-DeepFM-w/o-res | 0.949$\pm 0.010$ | 0.901$\pm 0.011$ | 0.882$\pm 0.011$ | 0.780$\pm 0.041$ | 0.828$\pm 0.024$ | 0.915$\pm 0.017$ | 0.759$\pm 0.031$ | 0.867$\pm 0.020$ |
| DSA-DeepFM-attn-rev | 0.957$\pm 0.030$ | 0.921$\pm 0.046$ | 0.885$\pm 0.065$ | 0.850$\pm 0.095$ | 0.867$\pm 0.081$ | 0.928$\pm 0.051$ | 0.811$\pm 0.113$ | 0.901$\pm 0.060$ |

The key difference between DSA-DeepFM-w/o-attn and DSA-DeepFM-w/o-res lies in their fusion approach. The former uses simple concatenation, while the latter employs a DSA mechanism (Figure 2). The results show that DSA-DeepFM-w/o-res significantly outperformed DSA-DeepFM-w/o-attn, with a mean AUC of 0.949 (±0.010) vs. 0.890 (±0.003). This highlights the attention mechanism’s capability for the integration of different input types and enhancement of feature representation.

DSA-DeepFM-w/o-res outperformed both DSA-DeepFM-cat (mean AUC: 0.897±0.004) and Attn-DeepFM-aux (mean AUC: 0.825±0.003), demonstrating that both input types offer unique advantages for prediction, and that their combination substantially improves performance. These results highlight the importance of our proposed fusion approach and residual structure in enhancing model performance.

We also examined the influence of the sequence of the two attention mechanisms by comparing DSA-DeepFM with DSA-DeepFM-attn-rev. DSA-DeepFM again demonstrated superior performance compared to DSA-DeepFM-attn-rev, with a mean AUC of 0.982 (±0.003) versus 0.957 (±0.030). Although reversing the attention order had a less significant impact than omitting the residual structure, it still yielded considerably better results than those obtained without the residual structure or when using only a single input type.

#### Drug ordering independence

We tested whether the order of drug pairs (i.e., drug A-drug B vs. drug B-drug A) affected the model’s predictions. Both combinations were included in the training and test sets during 5-fold cross-validation testing using the DrugCombDB dataset. Figure S1 shows the predicted probabilities for all test samples, with a Pearson correlation of 0.998 between the two drug orderings. This high correlation indicates that the model’s predictions are largely independent of drug ordering. The clustering of probabilities around (0, 0) and (1, 1) shows strong discriminatory ability of DSA-DeepFM.

#### Explainability: Visualization of embedding vectors at different stages of DSA-DeepFM

We visualized the embedding features of drug-drug-cell line samples from the test set at various stages using t-distributed Stochastic Neighbor Embedding (t-SNE) to evaluate the effectiveness of DSA-DeepFM (Figure 4, Section 1.4 of the Supplementary Document). Initially, the pre-training embedding layer showed no distinction between synergistic and antagonistic combinations. After training, the embedding layer improved class separation, especially for categorical data. Auxiliary data, while useful, required further processing, and the trained Feature Extraction module helped better differentiate the inputs. The fusion of categorical and auxiliary data through the attention mechanism resulted in clearer class separation, enhancing the model’s predictive power. By using the final prediction module, the model demonstrated strong classification ability, distinctly separating synergistic and antagonistic drug combinations. This analysis highlights the importance of feature fusion and the attention mechanism in improving model performance.

**Fig. 4.**
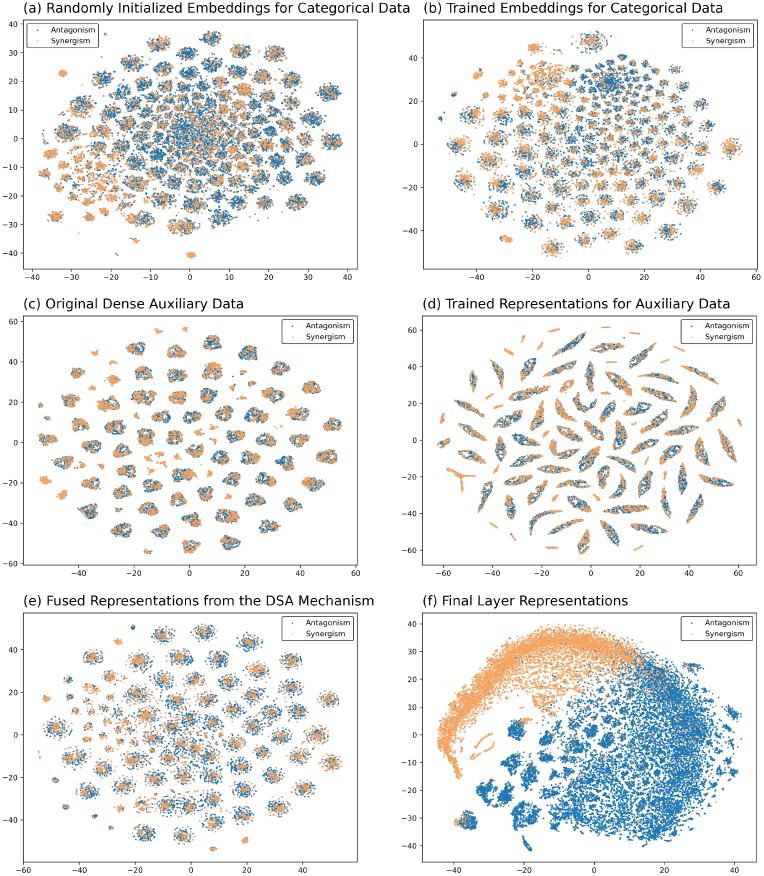
The t-SNE visualization of the representations of drug-drug-cell line triplets output at different stages of DSA-DeepFM. (a) Initialized embedding vectors from the embedding layer for categorical data before training. (b) Embedding vectors of the categorical data produced by the trained embedding layer. (c) Input auxiliary numerical data. (d) Representation vectors of the auxiliary data produced by the trained Feature Extraction module. (e) Fused representation vectors produced by the DSA mechanism that combines categorical and numerical vectors. (f) Final representation vectors used for prediction by the prediction module of the model.

### Validation Tests

Using five-fold cross-validation and leaving-cell line (or tissue)-out testing, we validated our method by comparing it with five traditional machine learning models (Random forest (RF), Support vector machine (SVM), Elastic net (EN), and Gradient Boosting machine (GBM) in the scikit-learn Python package and XGBoost in xgboost Python package) and four deep learning models (Multi-Layer Perceptron (MLP), MatchMaker [Kuru et al., 2021], DeepDDS [Wang et al., 2022a], and HypergraphSynergy [Liu et al., 2022]).

The average accuracy and variance from the five-fold cross-validation tests on the DrugCombDB and O’Neil datasets are summarized in Tables 3 and S1, respectively. From these tables, we derive the following findings regarding the prediction of the synergistic properties of drug combinations on cell lines.

**Table 3.** Performance of DSA-DeepFM and the nine compared models on the DrugCombDB dataset. The highest average accuracy values for the traditional machine learning methods and deep learning methods are highlighted in blue and purple, respectively.

| Model | AUC-ROC | ACC | Precision | Recall | F1 | AUC-PR | Kappa | BACC |
| --- | --- | --- | --- | --- | --- | --- | --- | --- |
| DSA-DeepFM | **0.982**±0.003 | **0.962** ±0.004 | **0.943**±0.006 | **0.932**±0.011 | **0.937** ±0.008 | **0.968**±0.005 | **0.910**±0.011 | **0.954**±0.006 |
| EN | 0.763±0.002 | 0.764±0.004 | 0.745±0.004 | 0.341±0.019 | 0.468±0.017 | 0.632±0.006 | 0.342±0.015 | 0.645±0.008 |
| GBM | 0.805±0.002 | 0.787±0.001 | 0.755±0.003 | 0.444±0.002 | 0.560±0.002 | 0.692±0.003 | 0.431±0.003 | 0.691±0.001 |
| RF | **0.888**±0.001 | **0.838**±0.002 | **0.831**±0.004 | **0.584**±0.006 | **0.686**±0.003 | **0.808**±0.002 | **0.581**±0.004 | **0.766**±0.003 |
| SVM | 0.771±0.001 | 0.741±0.001 | 0.740±0.005 | 0.227±0.001 | 0.347±0.005 | 0.616±0.002 | 0.239±0.004 | 0.596±0.002 |
| XGBoost | 0.832±0.002 | 0.800±0.001 | 0.783±0.004 | 0.473±0.001 | 0.589±0.001 | 0.728±0.003 | 0.467±0.001 | 0.708±0.001 |
| DeepDDS-GAT | 0.846±0.030 | 0.817±0.023 | 0.781±0.030 | 0.552±0.072 | 0.646±0.057 | 0.754±0.049 | 0.528±0.068 | 0.743±0.036 |
| DeepDDS-GCN | 0.857±0.029 | 0.818±0.023 | 0.781±0.019 | 0.555±0.083 | 0.647±0.060 | 0.764±0.048 | 0.530±0.071 | 0.744±0.040 |
| HypergraphSynergy | **0.905**±0.007 | **0.847** ±0.010 | 0.744±0.028 | **0.761** ±0.016 | **0.752** ±0.012 | **0.843** ±0.011 | **0.642** ±0.020 | **0.823** ±0.007 |
| MatchMaker | 0.886±0.006 | 0.844±0.003 | **0.789** ±0.006 | 0.666±0.013 | 0.722±0.008 | 0.810±0.007 | 0.615±0.009 | 0.794±0.005 |
| MLP | 0.858±0.009 | 0.819±0.009 | 0.723±0.027 | 0.657±0.016 | 0.688±0.011 | 0.774±0.014 | 0.560±0.018 | 0.773±0.007 |

- Among the five traditional machine learning methods, RF performs best on the DrugCombDB dataset, while XGBoost achieves the highest performance on the O’Neil dataset.
- HypergraphSynergy and MatchMaker are the top two compared deep learning models, with the former outperforming the latter by a small margin across all metrics except Precision.
- The accuracy improvements of our model over the other models on DrugCombDB are as least: 8% in AUC-ROC, 13% in ACC, 19% in Precision, 22% in Recall, 25% in F1, 15% in AUC-PR, 24% in Kappa, and 16% in BACC.
- The accuracy improvements of our model over the others on O’Neil dataset are at least: 5% in AUC-ROC, 13% in ACC, 17% in Precision, 10% in Recall, 14% in F1, 6% in AUC-PR, 31% in Kappa, and 13% in BACC.

In the more challenging leave-cell line-out and leave-tissue-out scenarios, we obtained mixed performance results (Tables 4 and 5). Our model significantly outperformed the alternatives in F1, Kappa, and BACC; however, it ranked second or third in the other five metrics.

**Table 4.** Performance of DSA-DeepFM and the nine compared models on the DrugCombDB Dataset in Leave-Cell-Line-Out testing. The highest average accuracy in each category is highlighted in blue or purple.

| Model | AUC-ROC | ACC | Precision | Recall | F1 | AUC-PR | Kappa | BACC |
| --- | --- | --- | --- | --- | --- | --- | --- | --- |
| DSA-DeepFM | **0.817**$\pm 0.011$ | **0.774**$\pm 0.010$ | **0.629**$\pm 0.032$ | **0.624**$\pm 0.036$ | **0.626**$\pm 0.026$ | **0.689**$\pm 0.031$ | **0.464**$\pm 0.024$ | **0.732**$\pm 0.012$ |
| EN | 0.672$\pm 0.060$ | 0.707$\pm 0.024$ | 0.528$\pm 0.059$ | 0.356$\pm 0.060$ | 0.423$\pm 0.048$ | 0.491$\pm 0.062$ | 0.237$\pm 0.055$ | 0.608$\pm 0.026$ |
| GBM | 0.722$\pm 0.046$ | 0.737$\pm 0.023$ | 0.594$\pm 0.052$ | 0.422$\pm 0.056$ | 0.492$\pm 0.052$ | 0.568$\pm 0.057$ | 0.323$\pm 0.060$ | 0.649$\pm 0.030$ |
| RF | 0.765$\pm 0.045$ | 0.763$\pm 0.023$ | **0.740**$\pm 0.055$ | 0.342$\pm 0.044$ | 0.466$\pm 0.042$ | 0.637$\pm 0.048$ | 0.340$\pm 0.047$ | 0.645$\pm 0.021$ |
| SVM | 0.610$\pm 0.028$ | 0.687$\pm 0.020$ | 0.466$\pm 0.053$ | 0.232$\pm 0.072$ | 0.305$\pm 0.075$ | 0.440$\pm 0.046$ | 0.138$\pm 0.060$ | 0.560$\pm 0.027$ |
| XGBoost | **0.788**$\pm 0.022$ | **0.764**$\pm 0.023$ | 0.627$\pm 0.044$ | **0.555**$\pm 0.046$ | **0.588**$\pm 0.035$ | **0.649**$\pm 0.040$ | **0.423**$\pm 0.048$ | **0.705**$\pm 0.024$ |
| DeepDDS-GAT | 0.791$\pm 0.022$ | 0.769$\pm 0.027$ | 0.663$\pm 0.033$ | 0.493$\pm 0.067$ | 0.563$\pm 0.044$ | 0.651$\pm 0.026$ | 0.411$\pm 0.053$ | 0.691$\pm 0.027$ |
| DeepDDS-GCN | **0.818**$\pm 0.021$ | **0.788**$\pm 0.014$ | **0.733**$\pm 0.051$ | 0.477$\pm 0.048$ | **0.576**$\pm 0.034$ | **0.690**$\pm 0.035$ | **0.444**$\pm 0.038$ | **0.701**$\pm 0.020$ |
| HypergraphSynergy | 0.729$\pm 0.035$ | 0.666$\pm 0.075$ | 0.478$\pm 0.049$ | **0.710**$\pm 0.098$ | 0.566$\pm 0.024$ | 0.586$\pm 0.041$ | 0.314$\pm 0.074$ | 0.677$\pm 0.032$ |
| MatchMaker | 0.750$\pm 0.034$ | 0.744$\pm 0.025$ | 0.589$\pm 0.050$ | 0.513$\pm 0.073$ | 0.546$\pm 0.056$ | 0.592$\pm 0.056$ | 0.369$\pm 0.068$ | 0.679$\pm 0.036$ |
| MLP | 0.690$\pm 0.015$ | 0.698$\pm 0.021$ | 0.506$\pm 0.037$ | 0.508$\pm 0.061$ | 0.504$\pm 0.023$ | 0.536$\pm 0.032$ | 0.287$\pm 0.014$ | 0.644$\pm 0.009$ |

**Table 5.** Performance of DSA-DeepFM and the nine compared models on the DrugCombDB Dataset in Leave-Tissue-Out testing. The highest average accuracy in each category is highlighted in blue or purple.

| Model | AUC-ROC | ACC | Precision | Recall | F1 | AUC-PR | Kappa | BACC |
| --- | --- | --- | --- | --- | --- | --- | --- | --- |
| DSA-DeepFM | **0.804**$\pm 0.100$ | **0.761**$\pm 0.073$ | **0.620**$\pm 0.139$ | **0.618**$\pm 0.201$ | **0.585**$\pm 0.143$ | **0.716**$\pm 0.124$ | **0.412**$\pm 0.117$ | **0.727**$\pm 0.089$ |
| EN | 0.688$\pm 0.127$ | 0.666$\pm 0.090$ | 0.525$\pm 0.189$ | 0.465$\pm 0.258$ | 0.421$\pm 0.146$ | 0.573$\pm 0.189$ | 0.194$\pm 0.105$ | 0.615$\pm 0.082$ |
| GBM | 0.732$\pm 0.089$ | 0.663$\pm 0.166$ | 0.559$\pm 0.189$ | 0.530$\pm 0.220$ | 0.487$\pm 0.148$ | 0.583$\pm 0.114$ | 0.265$\pm 0.132$ | 0.633$\pm 0.053$ |
| RF | 0.759$\pm 0.105$ | **0.719**$\pm 0.119$ | **0.685**$\pm 0.191$ | 0.407$\pm 0.211$ | 0.448$\pm 0.090$ | 0.639$\pm 0.098$ | 0.286$\pm 0.129$ | 0.649$\pm 0.092$ |
| SVM | 0.587$\pm 0.097$ | 0.690$\pm 0.126$ | 0.523$\pm 0.231$ | 0.288$\pm 0.242$ | 0.324$\pm 0.204$ | 0.455$\pm 0.193$ | 0.121$\pm 0.115$ | 0.555$\pm 0.055$ |
| XGBoost | **0.774**$\pm 0.118$ | 0.700$\pm 0.124$ | 0.590$\pm 0.202$ | **0.619**$\pm 0.184$ | **0.553**$\pm 0.144$ | **0.652**$\pm 0.109$ | **0.336**$\pm 0.162$ | **0.681**$\pm 0.067$ |
| DeepDDS-GAT | 0.798$\pm 0.080$ | 0.769$\pm 0.080$ | **0.649**$\pm 0.221$ | 0.437$\pm 0.173$ | 0.515$\pm 0.183$ | 0.702$\pm 0.120$ | 0.354$\pm 0.136$ | 0.666$\pm 0.064$ |
| DeepDDS-GCN | **0.818**$\pm 0.081$ | **0.774**$\pm 0.096$ | 0.617$\pm 0.298$ | 0.411$\pm 0.208$ | 0.492$\pm 0.243$ | **0.738**$\pm 0.113$ | **0.358**$\pm 0.180$ | 0.666$\pm 0.083$ |
| HypergraphSynergy | 0.742$\pm 0.038$ | 0.709$\pm 0.053$ | 0.514$\pm 0.150$ | **0.712**$\pm 0.146$ | **0.576**$\pm 0.143$ | 0.587$\pm 0.192$ | 0.329$\pm 0.104$ | **0.698**$\pm 0.054$ |
| MatchMaker | 0.740$\pm 0.089$ | 0.720$\pm 0.089$ | 0.571$\pm 0.166$ | 0.588$\pm 0.209$ | 0.539$\pm 0.132$ | 0.585$\pm 0.166$ | 0.319$\pm 0.136$ | 0.676$\pm 0.088$ |
| MLP | 0.654$\pm 0.090$ | 0.666$\pm 0.108$ | 0.492$\pm 0.157$ | 0.509$\pm 0.168$ | 0.470$\pm 0.107$ | 0.533$\pm 0.095$ | 0.221$\pm 0.114$ | 0.620$\pm 0.056$ |

### Discovery of Novel Synergistic Drug Combinations

We selected the ten cell lines with the highest frequencies in DrugCombDB and validated all possible drug combinations that were not included in the training set. We identified the top four drug pairs in terms of likelihood for the human colorectal carcinoma cell line HCT116 and the human ovarian cancer cell line SK-OV-3 (Figure 5). These findings are reinforced by other models (Table 6).

**Table 6.** Predicted outcomes from the nine compared models on the eight drug-drug-cell line combinations given in Figure 5, with a synergy cutoff score set at 0.6. Positive outcomes are shown in red. The three combinations validated by lab experiments are highlighted in bold in the first column.

| Drug Pair | Cell Line | EN | GBM | RF | SVM | XGBoost | DeepDDS-GAT | DeepDDS-GCN | Hypergraph | MatchMaker | MLP |
| --- | --- | --- | --- | --- | --- | --- | --- | --- | --- | --- | --- |
| Actinomycin D + Doxorubicin | HCT116 | 0.505 | **0.642** | **0.739** | 0.470 | **0.905** | 0.090 | 0.466 | 0.502 | **1.000** | **1.000** |
| **Etoposide + Geldanamycin** |  | **0.613** | **0.673** | **0.703** | 0.497 | **0.955** | 0.003 | 0.097 | 0.431 | **1.000** | **0.967** |
| **Etoposide + Vorinostat** |  | 0.461 | 0.459 | **0.624** | 0.350 | **0.910** | **0.624** | 0.414 | **0.929** | **1.000** | 0.402 |
| Imatinib + Mitoxantrone |  | 0.433 | 0.459 | 0.568 | 0.251 | **0.737** | 0.408 | 0.172 | **0.647** | 0.583 | **0.600** |
| Cabazitaxel + 5-Fluorouracil | SK-OV-3 | 0.549 | **0.636** | **0.629** | 0.265 | **0.919** | **0.612** | **0.651** | **0.749** | **0.996** | **1.000** |
| **Paclitaxel + Imatinib** |  | 0.328 | 0.364 | 0.423 | 0.251 | **0.731** | 0.304 | 0.336 | 0.461 | **1.000** | **0.993** |
| Paclitaxel + Vismodegib |  | 0.488 | 0.543 | 0.594 | 0.256 | **0.957** | 0.486 | 0.474 | **0.616** | **1.000** | **1.000** |
| Paclitaxel + Axitinib |  | 0.442 | 0.574 | 0.513 | 0.257 | **0.854** | 0.331 | 0.262 | 0.137 | **1.000** | **1.000** |

**Fig. 5.**
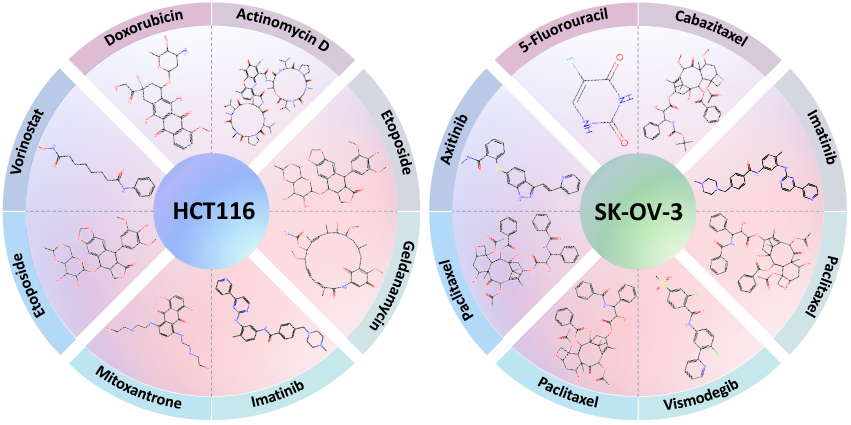
Top four predicted drug-drug combinations for the human colorectal carcinoma cell line HCT116 (left) and human ovarian cancer cell line SK-OV-3 (right).

Etoposide and Geldanamycin combination was identified to be synergistic in HCT116. This is consistent with that targeting both Topoisomerase II and HSP90 enzymes with the predicted drug combinations accelerates and enhances apoptosis, leading to a higher percentage of apoptotic cells compared to single-drug treatments in HCT116 [Barker et al., 2006].

Vorinostat and Etoposide are another drug pair predicted to have a synergistic effect in HCT116. This is consistent with the observation that Vorinostat as an HDAC inhibitor induces chromatin relaxation, thereby enhancing DNA accessibility to Etoposide, which amplifies Etoposide’s ability to induce DNA damage and promote cell death Alzoubi et al. [2016].

Furthermore, Gray et al. [2012] demonstrated that Vorinostat exhibited the strongest synergistic effect when combined with either Etoposide or Topotecan in several cell lines. Although their experiments were not done on HCT116, their explanation of drug interaction mechanisms indirectly supports the synergistic effect of the Vorinostat and Etoposide combination in this cell line. Additionally, Kumar et al. [2018] showed that Vorinostat, in combination with Etoposide, exhibited synergistic effects in HeLa cancer cell lines.

The third predicted synergistic drug combination is Paclitaxel and Imatinib in SK-OV-3. Mundhenke et al. [2008] showed that combining Paclitaxel with Imatinib at concentrations of 1 or 2 µM significantly reduced cell proliferation in SK-OV-3 and three other cell lines. When the concentration of Imatinib increased to 2 or 4 µM, the reduction in proliferation became even more pronounced.

Although the remaining five predicted drug combinations lack experimental validation, the involved drug pairs have demonstrated synergistic effects in other cell lines in previous studies. Below is some indirect evidence.

- Actinomycin D exhibits anti-tumor activity, and a study [Yang et al., 2024] on triple-negative breast cancer revealed that its combination with Doxorubicin significantly increased the apoptosis rate compared to either drug used alone.
- The combination of Mitoxantrone and Imatinib has been tested across various tumor types. Pinto et al. [2011] evaluated this combination in a prostate cancer model, demonstrating that it effectively reduces the required dosage of Mitoxantrone while achieving comparable growth inhibition. Furthermore, studies [Fruehauf et al., 2005, 2007] on chronic myeloid leukemia have shown that the Mitoxantrone and Imatinib combination is a well-tolerated treatment protocol.
- Ongoing clinical trials [Machiels et al., 2016, Camille et al., 2017] are investigating the dosages of cabazitaxel and 5-fluorouracil for treating locally advanced squamous cell carcinoma of the head and neck. Although these trials do not directly involve SK-OV-3, they provide valuable insights into the potential synergy of these drugs in this cell line.
- A preliminary study [Kozloff et al., 2012] involving 49 patients assessed the combination of axitinib and paclitaxel, administered twice daily over a 3-week treatment cycle. The results indicated that this combination is effective for advanced non-small cell lung cancer, further supporting the exploration of its synergistic effects in SK-OV-3.
- Lastly, research by Yeh et al. [2024] highlights that Vismodegib, a Hedgehog inhibitor, reduces Bax phosphorylation, thereby enhancing paclitaxel-induced cytotoxicity. This mechanism leads to mitochondrial damage and apoptosis in non-small cell lung cancer cells.

## Discussion and Conclusion

We have presented DSA-DeepFM for predicting the synergistic property of drug combinations. The model simultaneously processes both categorical information encoded through one-hot encoding and auxiliary numerical information on drugs and cell lines. The categorical input data leverage an embedding mechanism to extract features that are particularly relevant for classification, whereas the numerical data provide biologically useful information on drugs and cell lines.

The inclusion of the DSA mechanism enhances the model’s ability to learn shared patterns in both field and embedding spaces, resulting in features with stronger representational capabilities. The further integration of the deep FM module enables the model to capture both low-order and high-order feature interactions, thereby enhancing its discriminative capability.

The model exhibits high accuracy and stability in predictions compared to existing approaches on the DrugCombDB and O’Neil datasets. Identification of the eight novel drug combinations in HCT116 and SK-OV-3, supported partially by wet-lab studies in the literature on pharmacodynamics, highlights the potential in practical applicability of our model. Notably, none of the other compared deep learning models achieved the same prediction (Table 6).

However, further enhancements are necessary for the model to be fully applicable to patient-specific data. One key limitation, common to other deep learning models [Abbasi and Rousu, 2024], is that DSA-DeepFM performs less effectively in leave-one-out cross-validation scenarios. This reduction in performance is likely due to the embedding mechanism used in the model, which may struggle with limited data in such cases. To address this, incorporating mechanisms from general models like BERT could be a promising direction. BERT’s ability to generate contextually rich embeddings might help improve the model’s performance in these more challenging validation settings.

The success of DSA-DeepFM opens up opportunities to explore its application in other areas. One promising avenue is drug response prediction [Chen and Zhang, 2022, 2024], where the model’s ability to analyze multi-dimensional data and capture complex interactions could be leveraged to predict how cell lines or individual patients will respond to specific drug treatments. This expansion could contribute to more personalized and effective therapeutic strategies. Applications may also be possible in drug repurposing. Ongoing refinements and broader applications will be crucial for unlocking the model’s full potential in both research and clinical settings.

## Data and method availability

After preprocessing and integrating the aforementioned datasets, the DrugCombDB dataset used in this study comprises 48,985 synergistic and 112,106 antagonistic drug-drug-cell line entries, covering 1,412 distinct drugs and spanning 93 human cancer cell lines from 12 tissue types.

The O’Neil dataset in this study comprises 13,596 positive (synergistic) entries, and 15,378 negative (antagonistic) entries, covering 34 cancer cell lines and 38 anticancer drugs.

Additionally, we identified 4,634 genes associated with drug targets and KEGG pathways for the cell lines in both datasets.

These datasets are available on https://drive.google.com/drive/folders/1KQU7kKH-MFl2bJhsrLv1ZHxm35Nvhhsf?usp=drive_link.

Source code is available on https://github.com/gracygyx/DSA-DeepFM.

## Supplementary data

Supplementary Data are available at NAR Online

## Author contributions

YXG and YHS contributed to the conception and design of the experiment. YXG contributed to the draft of the paper. LXZ and JZ initiated and supervised the research. All authors contributed to the critical revision of the manuscript.

## Supplementary Materials

## 1. Model Training

### 1.1 Hyperparameter settings

We examine the influence of crucial hyperparameters on model performance, with particular attention to the embedding dimension of sparse categorical data in the FM Layer, the dimensionality of the two-layer network in the DNN component, the initial learning rate and the dropout rate during model training

#### Embedding dimension in the FM layer

The FM layer transforms high-dimensional sparse categorical data into a dense vector. We tested the dimension of the embedding vector of 2^*k*^ for *k* from 6 to 10. Figure 1a shows the impact of embedding dimension on model performance in AUC. The results indicate a consistent improvement in performance as the embedding dimension increases. Notably, when the embedding dimension reaches 2^9^ (i.e. 512), model’s performance stabilizes, with both the mean and variance of the AUC closely aligning with those observed at an embedding dimension of 2^10^ (i.e. 1024). Consequently, we selected 512 as the optimal embedding dimension.

#### Dimensions of the DNN component

The DNN component consists of a two-layer fully connected network designed to capture higher-order interactions among input features. We tested the following combinations of layer dimensions: 2^8^ × 2^8^ (i.e. 256 × 256), 2^8^ × 2^9^, 2^9^ × 2^9^, 2^9^ × 2^10^, and 2^10^ × 2^10^ to determine the optimal network size. Figure 3b illustrates the effect of these dimension combinations on the model’s performance, measured by AUC.

We began with a network size of 2^8^×2^8^. Doubling the neurons in the second layer led to an increase in the mean AUC and a reduction in variance. Next, when the neurons in the first layer were doubled, the mean AUC remained unchanged, but there was a slight increase in variance. Further doubling the neurons in the second layer resulted in a significant improvement in mean AUC and a notable decrease in variance. Finally, doubling the neurons in the first layer again left the mean AUC stable, with a slight decrease in variance, suggesting diminishing returns with further increases in network size.

To maintain a manageable number of model parameters, we decided not to expand the network size further. Given that the model’s performance was nearly identical for the 2^9^ × 2^10^ and 2^10^ × 2^10^ configurations, we ultimately adopted the 2^9^ × 2^10^ (i.e. 512 × 1024) fully connected network for the DNN component.

### 1.2. Learning rate

The model was trained using the Adam optimizer with learning rate decay. The initial learning rate was treated as a hyperparameter and tuned by selecting from the values 0.01, 0.001, and 0.0001. Figure 1c shows that reducing the learning rate from 0.01 to 0.001 led to a significant increase in the mean AUC and a decrease in variance. However, further decreasing the learning rate to 0.0001 resulted in a decline in the mean AUC. Based on these observations, we selected a learning rate of 0.001 for the model.

### 1.3 Dropout rate

The dropout rate is a crucial hyperparameter in machine learning models, particularly for preventing overfitting during training. In our experiments, we evaluated various dropout rates, including 0 (no dropout), 0.1, 0.2, 0.3, 0.4, and 0.5, to assess their impact on model performance. As shown in Figure 1d, increasing the dropout rate from 0 to 1 resulted in a decrease in the AUC value. Dropout rates between 0.1 and 0.3 showed a gradual increase in AUC, while a rate of 0.4 led to a decline. Notably, at a dropout rate of 0.5, the mean AUC reached its highest value, with the lowest variance in AUC scores. Overall, the dropout parameter had only a minimal impact on the model’s performance.

### 1.4 Visualization of embedding vectors for drug-drug-cell line triplets

To thoroughly analyze the effectiveness of our classification model, we visualized the embedding features of drug-drug-cell line samples in the test set at various stages using t-distributed Stochastic Neighbor Embedding (t-SNE) [Van der Maaten and Hinton, 2008]. The stages include:

- Embedding vectors produced by the embedding layer for the input categorical data before the layer was trained;
- Representation vectors from the trained embedding layer for the input categorical data;
- The input auxiliary data;
- Representation vectors from the trained two-layer network for the input auxiliary data;
- Fused representation from the trained attention mechanism, combining categorical and auxiliary vectors;
- Representation vectors produced by the trained prediction component on which prediction was made.

In Figure 4a, the scatter plot illustrates the embedded representations of the categorical inputs produced by the pre-training embedding layer. The drug-drug-cell line triplets with antagonistic effects (blue) and synergistic effects (yellow) exhibit similar distributions and are indistinguishable, reflecting the random initialization in the pre-training model.

In Figure 4b, the representations generated by the trained embedding layer for the categorical inputs are shown. Compared to the random distribution in Figure 4a, these representations demonstrate better separation between the two classes, with antagonistic samples (blue) becoming more compact. This suggests that the trained embedding layer enhances the separation between the two classes, forming distinct clusters.

Figure 4c shows the input auxiliary data. While the two types of data are somewhat distinguishable compared to Figure 4a, there remains a significant overlap between the positive and negative data entries. This indicates that the auxiliary data is useful, but requires further processing to fully realize its potential.

Figure 4d shows the representations from the auxiliary inputs produced by the trained two-layer network. Compared to the input auxiliary data in Figure 4c, the representations are more dispersed, indicating greater differentiation between data points. However, distinguishing between the two classes remains challenging at this stage. A comparison of Figure 4c and Figure 4d shows that categorical inputs are more tightly clustered, with well-defined class centers, while numerical inputs are more scattered. This suggests that fusing both input types could enhance classification performance.

Figure 4e visualizes the representations of the fused categorical and auxiliary embeddings processed through the trained attention mechanism. Compared to Figure 4c, the data points in Figure 4e are more dispersed, with distinct class centers, whereas the class centers in Figure 4c were less pronounced. This broader dispersion of samples, along with clearer class centers, demonstrates the effectiveness of our proposed fusion approach in enhancing the model’s predictive capabilities, as expected from leveraging the attention mechanism.

Figure 4f illustrates the representations from the final layer after training. At the final stage, the two classes of test samples are distinctly separated in the two-dimensional space, demonstrating the model’s strong ability to predict the synergistic property.

This analysis demonstrates that the proposed fusion strategy, attention mechanism, and overall model each play a crucial role in refining feature representation and enhancing the model’s classification performance. The transformation of the data points becomes increasingly effective, leading to improved accuracy in distinguishing between synergistic and antagonistic drug combinations.

## 2. Metrics for Measuring Prediction Accuracy

We consider drug combination prediction as a binary classification problem. Define the true positive rate (TPR) to be the proportion of actual positives that are correctly identified by the model, that is

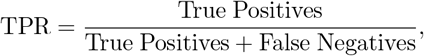

and define the false positive rate (FPR) as:

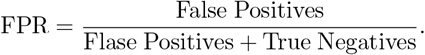

Eight metrics are used to evaluate different models, including:

- **Area under the Receiver Operating Characteristic curve** (AUC-ROC): The ROC curve plots the TPR (y-axis) against the FPR (x-axis) at *n* different threshold values *α*_1_, *α*_2_, · · ·, *α*_*n*_. By adjusting the threshold for classification, the ROC curve shows the trade-offs between sensitivity (true positives) and specificity (false positives) for a binary classification problem. AUC-ROC is defined as:

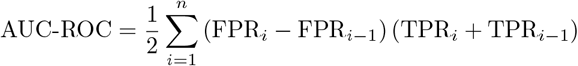

where TPR_*i*_ and FPR_*i*_ are the TPR and FPR at the threshold *α*_*i*_.

- **Recall**: the TPR is also called the recall.
- **Precision**: It is the ratio of True Positives to the sum of True Positives and False Positives.
- **Area under the precision-recall curve** (AUC-PR):

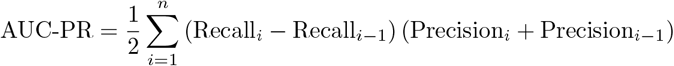

where Recall_*i*_ and Precision_*i*_ are the Recall and Precision at the *i*-th threshold value.

- **Accuracy** (ACC): It is the ratio of correctly predicted instances to the total instances, that is

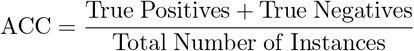

- **F1 score**:

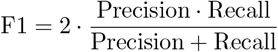

- **Cohen’s Kappa** (Kappa):

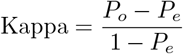

where *P*_*o*_ is the identical prediction rate between the two classifiers and *P*_*e*_ is the proportion of agreement that would be expected by chance alone, based on the distribution of categories assigned by each classifier.

- **Balanced accuracy** (BACC): It is the half of the sum of TPR and TNR.

## 3. Deep Learning Models Examined During Validation Testing

We compared the proposed model with the following deep learning methods during validation testing.

- MatchMaker (Kuru et al., IEEE-ACM TCBB, 2021): It comprises two parts: the drug-specific subnetworks and the synergy prediction subnetwork. The former contains two parallel sub-networks that learn the representations of each drug in a specific cell line. The latter consists of fully connected layers to predict the synergistic effects of drugs.
- DeepDDS (Wang et al., Briefings in Bioinformatics, 2022): It employs graph networks and attention mechanisms to predict the effect of drug combinations. Specifically, it uses an MLP to extract features from cell line gene expression data and uses a graph attention network or a graph convolution network to extract drug features based on molecular graphs. Finally, the embedding vectors of drug-pair-cell lines are then concatenated to predict the property of the combination. Depending on the model used for drug feature extraction, DeepDDS can be referred to as DeepDDS-GAT or DeepDDS-GCN.
- HypergraphSynergy (Liu et al., Bioinformatics,2022): It models drug-drug-cell line combinations as a hypergraph, where the nodes represent drugs and cell lines, and the hyperedges represent drug-drug-cell line interactions. By leveraging a hypergraph neural network, HypergraphSynergy generates embedding vectors for both drugs and cell lines, which are then combined to predict the synergistic properties of drug combinations.

Note that MatchMaker was originally designed to solve drug combination prediction as a regression task. To adapt them for classification, we replaced the activation function in the final layer with a sigmoid function and substituted the loss function with cross-entropy loss.

**Figure S1.**
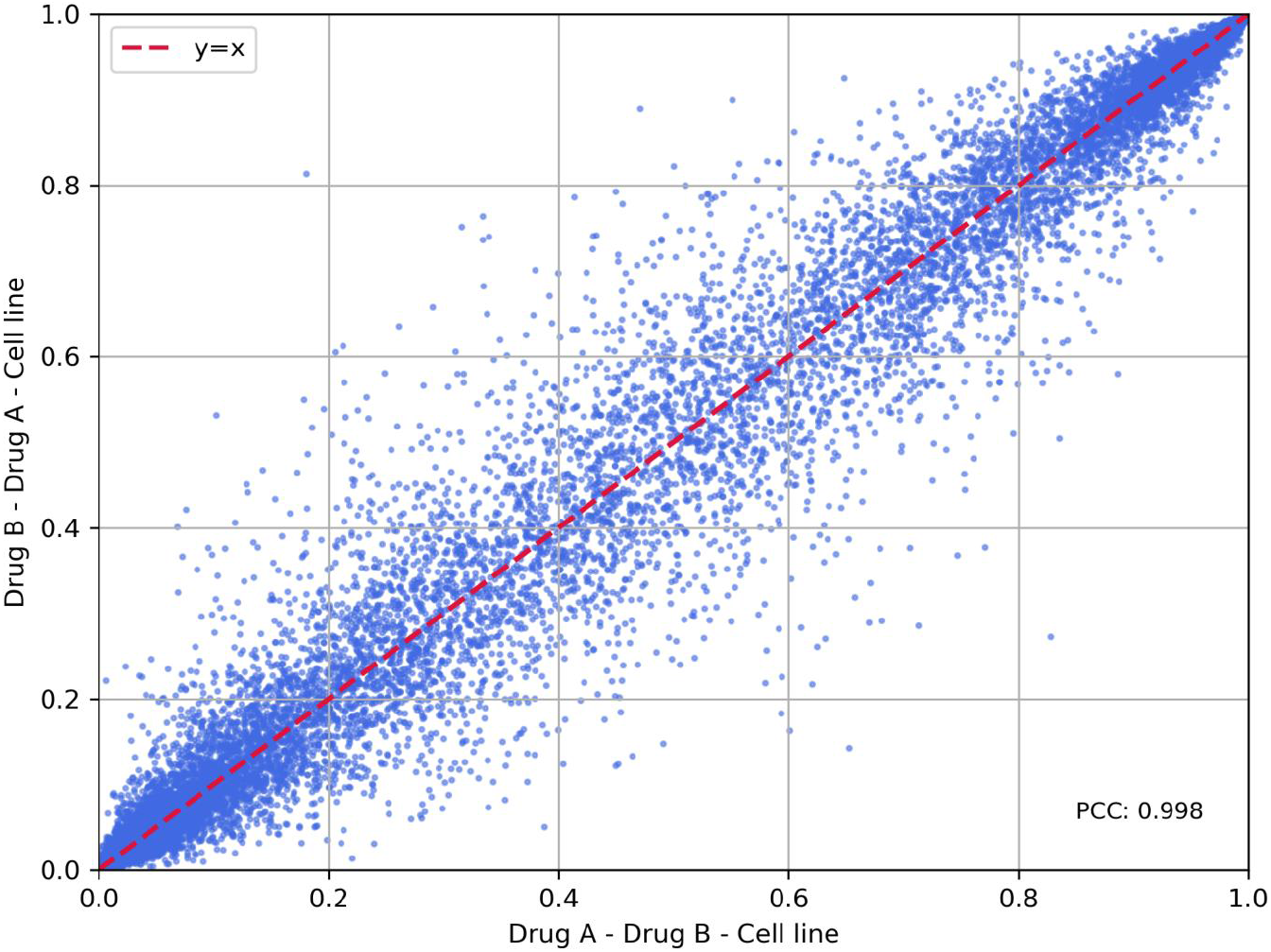
Scatter plot of predicted probabilities under different drug input orders.

**Table S1.**
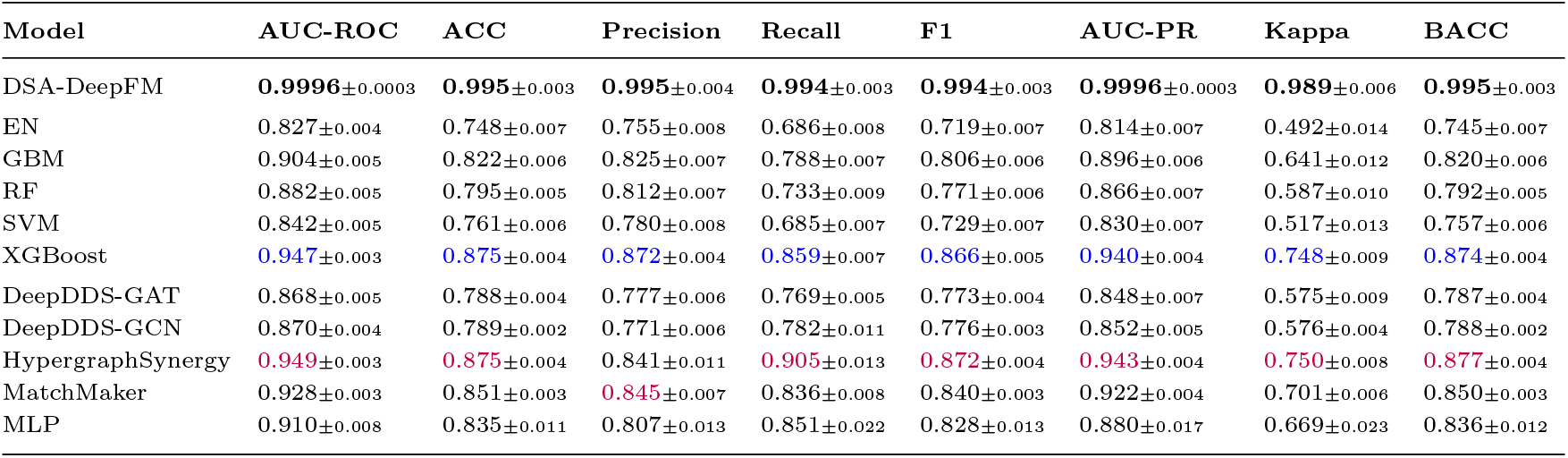
Performance of DSA-DeepFM and the nine compared models on the O’Neil dataset. The highest average accuracy values for the traditional machine learning methods and deep learning methods are highlighted in blue and purple, respectively.

| Model | AUC-ROC | ACC | Precision | Recall | F1 | AUC-PR | Kappa | BACC |
| --- | --- | --- | --- | --- | --- | --- | --- | --- |
| DSA-DeepFM | **0.9996**$\pm 0.0003$ | **0.995**$\pm 0.003$ | **0.995**$\pm 0.004$ | **0.994**$\pm 0.003$ | **0.994**$\pm 0.003$ | **0.9996**$\pm 0.0003$ | **0.989**$\pm 0.006$ | **0.995**$\pm 0.003$ |
| EN | 0.827$\pm 0.004$ | 0.748$\pm 0.007$ | 0.755$\pm 0.008$ | 0.686$\pm 0.008$ | 0.719$\pm 0.007$ | 0.814$\pm 0.007$ | 0.492$\pm 0.014$ | 0.745$\pm 0.007$ |
| GBM | 0.904$\pm 0.005$ | 0.822$\pm 0.006$ | 0.825$\pm 0.007$ | 0.788$\pm 0.007$ | 0.806$\pm 0.006$ | 0.896$\pm 0.006$ | 0.641$\pm 0.012$ | 0.820$\pm 0.006$ |
| RF | 0.882$\pm 0.005$ | 0.795$\pm 0.005$ | 0.812$\pm 0.007$ | 0.733$\pm 0.009$ | 0.771$\pm 0.006$ | 0.866$\pm 0.007$ | 0.587$\pm 0.010$ | 0.792$\pm 0.005$ |
| SVM | 0.842$\pm 0.005$ | 0.761$\pm 0.006$ | 0.780$\pm 0.008$ | 0.685$\pm 0.007$ | 0.729$\pm 0.007$ | 0.830$\pm 0.007$ | 0.517$\pm 0.013$ | 0.757$\pm 0.006$ |
| XGBoost | 0.947$\pm 0.003$ | 0.875$\pm 0.004$ | 0.872$\pm 0.004$ | 0.859$\pm 0.007$ | 0.866$\pm 0.005$ | 0.940$\pm 0.004$ | 0.748$\pm 0.009$ | 0.874$\pm 0.004$ |
| DeepDDS-GAT | 0.868$\pm 0.005$ | 0.788$\pm 0.004$ | 0.777$\pm 0.006$ | 0.769$\pm 0.005$ | 0.773$\pm 0.004$ | 0.848$\pm 0.007$ | 0.575$\pm 0.009$ | 0.787$\pm 0.004$ |
| DeepDDS-GCN | 0.870$\pm 0.004$ | 0.789$\pm 0.002$ | 0.771$\pm 0.006$ | 0.782$\pm 0.011$ | 0.776$\pm 0.003$ | 0.852$\pm 0.005$ | 0.576$\pm 0.004$ | 0.788$\pm 0.002$ |
| HypergraphSynergy | 0.949$\pm 0.003$ | 0.875$\pm 0.004$ | 0.841$\pm 0.011$ | 0.905$\pm 0.013$ | 0.872$\pm 0.004$ | 0.943$\pm 0.004$ | 0.750$\pm 0.008$ | 0.877$\pm 0.004$ |
| MatchMaker | 0.928$\pm 0.003$ | 0.851$\pm 0.003$ | 0.845$\pm 0.007$ | 0.836$\pm 0.008$ | 0.840$\pm 0.003$ | 0.922$\pm 0.004$ | 0.701$\pm 0.006$ | 0.850$\pm 0.003$ |
| MLP | 0.910$\pm 0.008$ | 0.835$\pm 0.011$ | 0.807$\pm 0.013$ | 0.851$\pm 0.022$ | 0.828$\pm 0.013$ | 0.880$\pm 0.017$ | 0.669$\pm 0.023$ | 0.836$\pm 0.012$ |

